# Epigenetic and transcriptional plasticity drive inflammation in Cystic Fibrosis

**DOI:** 10.1101/2022.10.12.511886

**Authors:** Adam M. Dinan, Odiri Eneje, Karen P. Brown, Frances Burden, Mary Morse, Rab K. Prinjha, Mattia Frontini, R. Andres Floto

## Abstract

The relative contributions of acute and chronic inflammatory mechanisms in diseases characterised by persistent bacterial infection are unclear, despite important consequences for the development of novel anti-inflammatory therapies. Here, we examined longitudinal transcriptional and epigenetic changes in patients with Cystic Fibrosis (CF), a genetic condition characterised by persistent bacterial lung infections that drive progressive inflammatory lung damage. We find that sudden clinical deteriorations in lung health, termed Acute Pulmonary Exacerbations (APEs), are linked to innate immune signalling driven by bacterial components, and accompanied by changes in macrophage function. Treatment of patients with intravenous antibiotics results in rapid modification of myeloid cell gene expression and epigenetic state, towards that of healthy volunteers, suggesting that repeated acute inflammatory episodes play an important role in CF inflammatory lung damage.

## Introduction

Cystic fibrosis (CF) is a genetic disease associated with dysfunction of the CF transmembrane conductance regulator (CFTR). Impaired mucociliary clearance leads to the colonisation of the lungs by bacterial pathogens, and a pattern of inflammation and infection is established. Cytotoxic products released by neutrophils to kill bacteria damage tissues and are associated with lung function decline (**1**).

Inflammation in CF has generally been regarded as chronic and continuous in nature (**1; 2**). However, there is also clinical evidence that cyclical processes are involved. CF patients frequently undergo episodes of rapid clinical deterioration (APEs) (**3**) that require prolonged (usually 14 days) treatment with intravenous antibiotics. It is not clear to what extent inflammation during such periods diverges from the baseline observed when the patient is clinically stable. CF therefore provides a unique opportunity to understand the temporal dynamics of inflammation in humans in the context of long-term bacterial infection.

## Results and Discussion

We collected whole blood samples from adult CF patients (*n* = 13) infected with *Pseudomonas aeruginosa* (PsA) at 3 timepoints: at an APE onset, at the end of intravenous antibiotic treatment, and when returned to stable clinical baseline (at least 30 days after the APE onset; **Fig. 1a**; **Dataset 01**). Analyses were performed on peripheral blood myeloid cells, due to the technical difficulty in isolating lung myeloid cells. Treatment with intravenous antibiotics was associated with resolution of systemic inflammation and an improvement in lung function (**Fig. 1b**). The study group was broadly representative of the clinical characteristics of the entire cohort attending the adult CF centre from which they were recruited (**Fig. 1c**).

**Figure 1.**
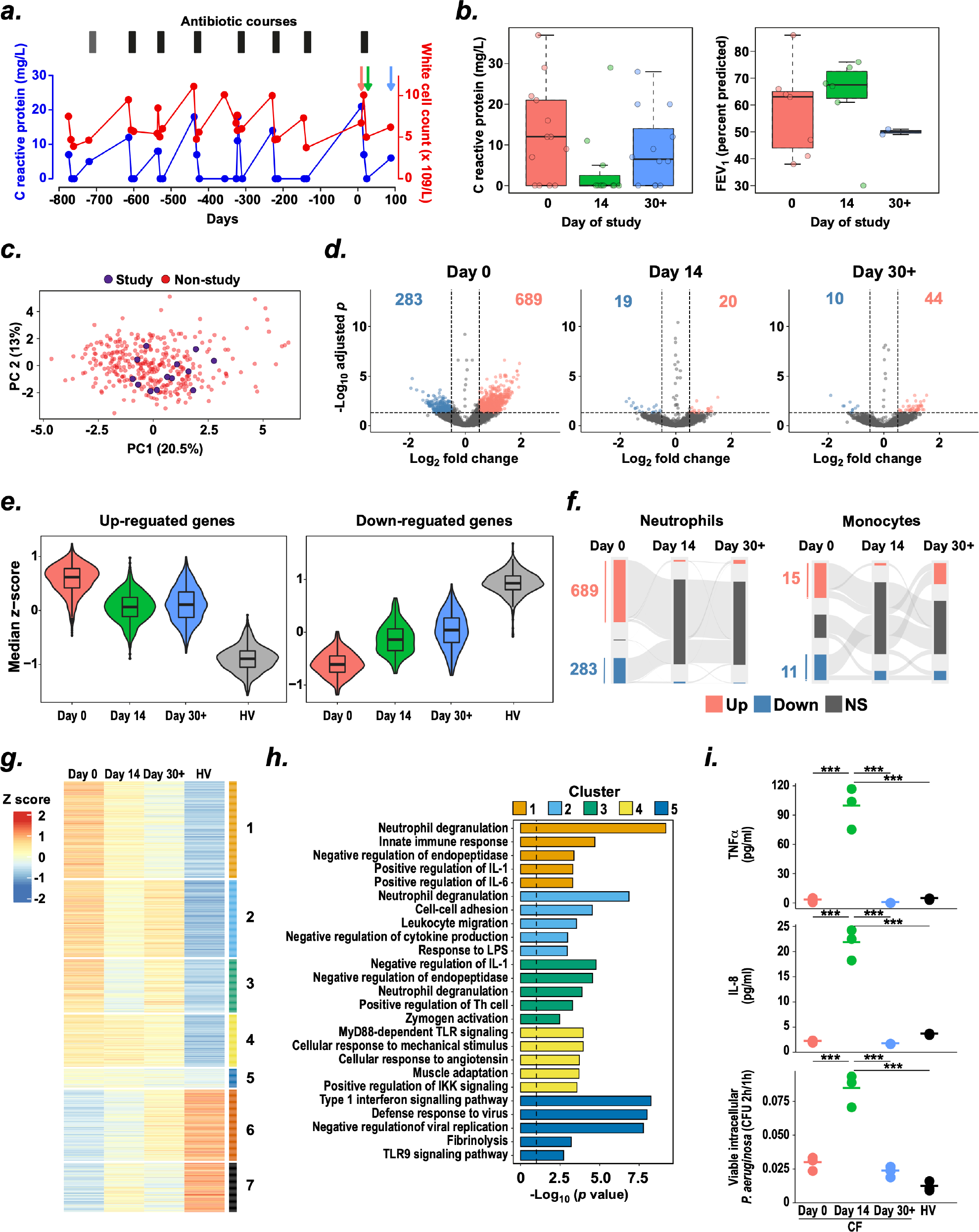
(**a**) Longitudinal fluctuation in c-reactive protein (CRP) and total white blood cell (WBC) count over a three-year period in a patient recruited for this study. Red arrow indicates Day 0; green arrow indicates day 14; blue arrow indicates Day 30+. (**b**) Levels of CRP and percent predicted forced expiratory volume in one second (FEV_1_ % predicted) for study patients at each sampling day. (**c**) Principal component analysis of clinical data set for 354 CF patients. Patients selected for inclusion in this study are indicated in purple. (**d**) Volcano plots show differentially expressed (DE) genes in CF neutrophils relative to HV at the indicated time points. (**e**) Median expression levels of DE genes by time point and study group. (**f**) Gene expression dynamics in neutrophils and monocytes. Parallel set diagrams show the changes in numbers of DE genes over sampling days (**g**) Unsupervised hierarchical clustering of neutrophil DE genes. Median z-scores per sample group are plotted for each gene. Gene clusters are indicated with coloured bars above heatmap. (**h**) GO term functional enrichment of neutrophil DE genes by cluster. Dashed line represents adjusted *P* value = 0.05. (**i**) Cytokine release and intracellular bacterial killing by monocyte-derived macrophages. Assays were performed at 4 hours following exposure of cells to PsA. Data from an example CF patient is shown. *** Adjusted *P* < 0.001, one-way ANOVA with Tukey’s post-hoc test.

Gene expression of peripheral neutrophils and monocytes isolated from CF patients on each sampling day was directly compared with samples from age- and sex-matched healthy volunteers (HV; *n* = 8; **Fig. 1d**). We observed a marked decrease in differential gene expression between CF and HV across the time series, with more differentially expressed (DE) genes at the onset of exacerbation (day 0), than on subsequent sampling days (days 14 and 30+) (**Fig 1d; Datasets 02 & 03**). The median expression level of DE genes across patients was responsive to antibiotic treatment, indicating that the effect was distributed across individuals and was not driven by subset of the study patients (**Fig 1e**).

Given the clear temporal dynamics of DE genes (**Fig 1f**), we used unsupervised hierarchical clustering to identify 7 gene clusters in neutrophils (**Fig. 1g**). Clusters of upregulated genes in CF were enriched for processes involved in innate immune response (**Fig. 1h**), while downregulated clusters showed no immune-specific enrichment.

We also found differences in the functional properties of monocyte-derived macrophages between day 0, day 14, and day 30+, with decreased intracellular killing and greater inflammatory cytokine production observed in Day 14 cells (**Fig. 1i**).

We next assessed changes at the epigenetic level, in both cell types, through genome-wide profiling of H3K27ac by chromatin immunoprecipitation (ChIP-seq). Differentially acetylated regions (DAR) were identified for each time point by direct comparison with HV samples (**Datasets 04 & 05**). Again, fewest DAR were identified in both neutrophils and monocytes at Day 14, indicating that IV treatment causes a reduction of the effects observed at day 0.

Two genes (*TLR5* and *MAP3K20*) had coordinate changes in transcription and H3K27ac at all time points in neutrophils (**Fig. 2a; Dataset 06**), while no genes had coordinate changes in monocytes (**Dataset 07**). In total, four DARs were assigned to *TLR5* (see **SI Appendix, Materials and Methods; Fig. 2a**), spanning the promoter region and a distal site located 57 kb upstream that has not previously been associated with *TLR5* expression (**Fig. 2b**).

**Figure 2.**
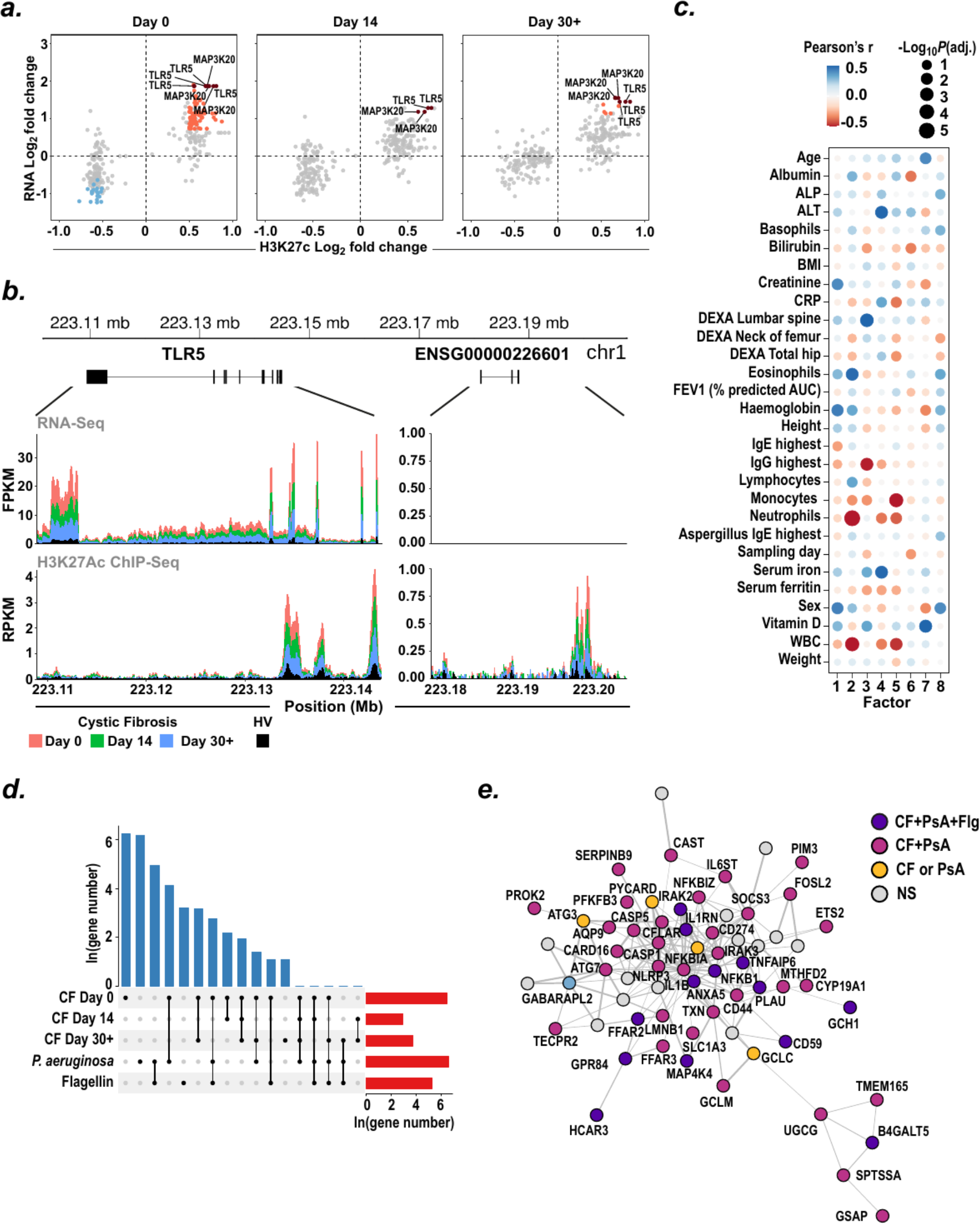
(**a**) RNA-Seq and ChIP-Seq log2 fold change values by sampling day. All differentially acetylated H3K27Ac peaks assigned to genes are shown as dots. Red dots, increased in CF relative to HV; blue dots, decreased in CF relative to HV; grey dots, no significant difference. (**b**) RNA-Seq and H3K27Ac ChIP-Seq coverage of the *TLR5* gene (left panel) and distal region of acetylation (right panel). The total coverage is shown as the median fragments per kilobase per million (FPKM) for RNA-Seq, and median reads per kilobase per million (RPKM) for ChIP-Seq, in 40 bp bins. (**c**) Correlation of clinical data with MOFA factors. Abbreviations: ALP, alkaline phosphatase; ALT, alanine transaminase; Asp, Aspergillus; AUC, area under curve; BMI, body mass index; CRP, C-reactive protein; FEV1 % pred., forced expiratory volume in one second (FEV1) percent of predicted; IgE, immunoglobulin E; IgG, immunoglobulin G; RAST, radioallergosorbent; WBC, total white blood cell count. (**d**) UpSet plot shows number of neutrophil upregulated genes in *P. aeruginosa*-infected HV neutrophils, flagellin-exposed HV neutrophils, and in CF at Day 0, Day 14, and Day 30+. (**e**) Functional network of genes commonly upregulated in *P. aeruginosa*-infected HV neutrophils (PsA) and CF neutrophils, constructed using interactions from the STRING database. A subset of genes in this network are also upregulated upon flagellin (flg) exposure. NS, not significantly differentially expressed.

We integrated all data layers for neutrophils and monocytes using a multi-omics factor analysis (MOFA) (**4**) and identified eight latent factors accounting for 30-60% of the variance across data types (**Figure 2c**). Several clinical variables were significantly correlated with latent MOFA factors (**Figure 2c**), indicating that phenotypic variation among CF patients is reflected at gene expression level and in the epigenetics of innate immune cells.

As all patients in this study were infected with PsA, we sought to assess the extent to which direct detection of PsA by myeloid cells was responsible for the observed changes. Neutrophils from three healthy volunteers were exposed to PsA or purified PsA flagellin (a known ligand for TLR5 (**5**)) and their transcriptional response was determined using RNA-seq.

As expected, the majority (86%) of the genes responsive to flagellin were also differentially expressed in PsA-exposed cells (*P* = 8.5 × 10^-207^) (**Fig. 2d**). PsA-upregulated genes overlapped with genes overexpressed significantly in CF neutrophils at day 0 (86 genes, *P* = 5.0 × 10^-13^) and at day 30+ (7 genes, *P* = 0.01), but not at day 14 (2 genes, *P* = 0.31; **Fig. 2**; **Dataset 08**).

Genes upregulated in PsA-exposed neutrophils and CF patient neutrophils formed a highly connected functional network (STRING (**6**) protein-protein interaction enrichment *P* value < 1 × 10^-16^). The largest connected component of this network included a subset of genes directly responsive to flagellin exposure (**Fig. 2e**), implicating this bacterial product as an important inflammatory driver. Moreover, 23 of the 88 proteins encoded by the genes in the network are responsive to one or more approved drugs (**Dataset 09**) and a further 6 to therapeutic compounds not yet approved (**Dataset 09**).

In summary, we have shown that the epigenetic, transcriptional, and functional properties of CF patient myeloid cells fluctuate temporally during the infection cycle. Additional work is required to determine whether or not the inflammatory program changes between periods of disease exacerbation. Peripheral myeloid cells have a short half-life, and the lack of persistence of epigenetic changes between sampling dates indicates that the bone marrow progenitors of these cells are not permanently modified during the infection cycle. Although we cannot exclude the possibility that antibiotic treatment directly modulates gene expression in these cells, the reduction in the expression of pro-inflammatory genes persists between Days 14 and 30+, indicating that it is more probably an indirect result of the effect of reducing bacterial burden in the lung.

We observed increased TNF production and decreased bacterial killing from macrophages isolated at Day 14 relative to the other sampling days. This may indicate that macrophages at Day 14 are sensitised to the presence of bacteria, due to the reduction in the infection burden caused by the administration of intravenous antibiotics. However, further work is necessary to establish the exact basis of such fluctuations. We also defined a functionally connected network of genes, which is directly responsive to PsA exposure. Targeting this network could therefore offer a rational strategy for reducing inflammation-associated pathology in CF.

## Materials and Methods

Patients with Cystic Fibrosis (CF) were enrolled from Royal Papworth Hospital, UK, after written informed consent (ethical approval REC 19/EE/0241). Peripheral blood samples were collected and processed using established protocols from the BLUEPRINT consortium (**7**). For the cytokine production assay, experiments were conducted using previously described methods (**8**). A detailed description of all Materials and Methods is available in **SI Appendix, Materials and Methods**.

## Data, Materials, and Software Availability

Sequence data have been deposited at the European Genome-phenome Archive (EGA) under accession number EGAS00001006421. Data files and code used for clustering and functional enrichment of differentially expressed genes can be accessed at https://doi.org/10.5281/zenodo.7113651.

## Acknowledgements

This work was supported by: University of Cambridge/GSK Varsity research award; UK Cystic Fibrosis Trust; British Heart Foundation; and The Wellcome Trust. The authors would like to thank Victoria Higgins for administrative support and Uche Agwo for contracting support.

## References

1. M. Cohen-Cymberknoh, E. Kerem, T. Ferkol, A. Elizur, Airway inflammation in cystic fibrosis: molecular mechanisms and clinical implications. Thorax 68, 1157–1162 (2013).

2. V. De Rose, Mechanisms and markers of airway inflammation in cystic fibrosis. Eur. Respir. J. 19, 333–340 (2002).

3. C. H. Goss, J. L. Burns, Exacerbations in cystic fibrosis. 1: Epidemiology and pathogenesis. Thorax 62, 360–367 (2007).

4. R. Argelaguet, et al., MOFA+: a statistical framework for comprehensive integration of multi-modal single-cell data. Genome Biol. 21, 111 (2020).

5. S.-I. Yoon, et al., Structural basis of TLR5-flagellin recognition and signaling. Science 335, 859–864 (2012).

6. D. Szklarczyk, et al., The STRING database in 2021: customizable protein-protein networks, and functional characterization of user-uploaded gene/measurement sets. Nucleic Acids Res. 49, D605–D612 (2021).

7. D. Adams, et al., BLUEPRINT to decode the epigenetic signature written in blood. Nat. Biotechnol. 30, 224–226 (2012).

8. L. Hepburn, et al., Innate immunity. A Spaetzle-like role for nerve growth factor β in vertebrate immunity to Staphylococcus aureus. Science 346, 641–646 (2014).

